# Insulin Does Not Augment *In Vitro* Tumor Growth Under A Hyperglycemia-Mimicking Milieu And In A Calorie Restriction-Resembling Manner

**DOI:** 10.1101/169664

**Authors:** Tao Liao, Xiao-Hui Li, Yan-Ping Chen, Li-Li Tan, Ji-Da Zhang, Xin-An Huang, Qin Xu, Sui-Qing Huang, Chang-Qing Li, Qing-Ping Zeng

## Abstract

**Background:** Whether insulin enhances or represses tumor cell proliferation remains debating and inconclusive although epidemiological data indicated insulin use raises a risk of cancer incidence in patients with diabetes mellitus (DM).

**Methodology/Principle Findings:** We cultured rat pituitary adenoma cells in a high-glucose medium to simulate hyperglycemia occurring in DM patients. Upon incubation with or without insulin, repressed tumor cell proliferation and downregulated tumor marker expression occur accompanying with mitigated oxidative stress and compromised apoptosis. Mechanistically, insulin resistance-abrogated glucose uptake was suggested to create an intracellular low-glucose milieu, leading to cellular starvation resembling calorie restriction (CR). While downregulation of insulin-like growth factor 1 (IGF-1) occurring in CR was validated, oncogene downregulation and tumor suppressor gene upregulation seen in CR was also replicated by *NOS2* knockdown.

**Conclusions/Significance:** Cellular starvation can exert CR-like anti-tumor effects regardless of insulin presence or absence.

## Introduction

Until 2015, approximately 415 million adults aged 20-79 have been estimated to live with diabetes mellitus (DM) worldwide, including 193 million who are undiagnosed [1]. From 2012-2015, 1.5 to 5.0 million deaths annually resulted from DM [2]. As an encouraging news for combating DM, the long-acting human insulin analogue insulin glargine with a relatively durable (18-26 h) hypoglycemic effect has been developed and widely marketed with a trade name of Lantus or Toujeo [3]. However, this improved human insulin derivative unfortunately has irritated a public concern of safety because the independent epidemiological investigations have associated a raised cancer risk with insulin use in DM patients [4-7]. Although the latest conclusions from meta-analyses of case reports has excluded a likelihood of insulin glargine in tumor induction [8,9], evidence supporting its potential risk in promoting cancer onset is continuing to emerge from the recent clinical surveys [10,11].

It seems a tremendous challenge to figure out why the previous epidemiological reports regarding insulin glargine’s effects on tumorigenesis is inconsistent before the *bona fide* etiological initiators of DM and cancer become identified [12], but it should be possible to clarify whether insulin enhances or represses tumor growth in DM patients. For this purpose, we assessed a putative correlation of insulin use with tumor growth in the *in vitro* cultured rat pituitary adenoma cell line GH3. Pituitary adenomas are common intracranial tumors accounting for 25% of brain tumors, which are generally benign but can cause significant morbidity via deregulated hormone production. Current strategies of therapy include medication, surgery, and radiotherapy [13]. However, there remains a substantial proportion of patients whose tumors are refractory to surgical and medical therapies. Therefore, other medical options attacking the tractable pituitary adenomas are needed urgently by targeting their aberrant metabolic patterns such as enhanced PI3K-Akt-mTOR signaling [14, 15].

To preliminarily investigate whether insulin would enhance *in vitro* pituitary adenoma proliferation, we cultured GH3 cells in a high-glucose (17.49 mM) or a higher-glucose (30 mM) medium supplemented with or without insulin (10^−9^ - 10^−7^ M). The glucose concentrations of 17.49 mM and 30 mM were chosen to mimic hyperglycemia seen in DM patients because 25 mM glucose in the medium is equal to hyperglycemia, while 5.5 mM glucose in the medium corresponds to normoglycemia [16]. It was previously indicated that under a sustained hyperglycemic condition, glucose uptake is impeded due to downregulation of glucose transporter 1 (GLUT1) [16], resembling insulin resistance seen in type 2 diabetes mellitus (T2DM). In practical, 10, 15, and 25 mM glucose were used to induce insulin resistance in 3T3-L1 adipocytes [17]. Similarly, 10^−7^ M insulin was also used to induce insulin resistance in HepG2 hepatoma cells [18], so we chose the insulin dose range from 10^−9^ M to 10^−7^ M.

Insulin resistance would prohibit glucose uptake and hence create an intracellular low glucose milieu, so-called cellular starvation, which mimics calorie restriction (CR) to reverse the process of cancer progression, invasion, and metastasis [19, 20].

Because CR was validated to block NF-kB responsive *NOS2* that encodes inducible nitric oxide synthase (iNOS) [21], we exploited *NOS2* knockdown to reveal a possible mechanism underlying cellular starvation exerting anti-tumor effects in a CR-like manner. If this is true, it should predicted that insulin resistance-induced hyperglycemia might decrease rather than increase the risk of cancer progression in T2DM-combined cancer patients upon insulin administration. Indeed, a conclusion that insulin does not enhance tumor cell proliferation in a high-glucose circumstance has been drawn from the present study.

Nevertheless, insulin use in type 1 diabetes mellitus (T1DM) patients with insulin deficiency or pro-diabetes individuals with insulin sensitivity might increase the cancer risk because of an intracellular high glucose milieu, which can activate the PI3K-Akt-mTOR signaling pathway to promote cell division [19, 20]. It should be essential, therefore, to accurately evaluate the cancer risk upon insulin use by distinguishing insulin resistance from insulin sensitivity. Our prospective anticipation regarding a causal link of insulin use to cancer incidence and progression merits further *in vivo* studies.

## Results

### Insulin-combined high-level glucose represses tumor cell proliferation and diminishes growth hormone secretion

Upon incubation with 10^−7^ M insulin for 24 or 48 h, GH3 cells exhibit a slightly faster growth rate than untreated GH3 cells albeit without statistical significance. However, when measured on 72 h, 10^−7^ M insulin remarkably delays GH3 cell growth with significant difference from untreated GH3 cells (Figure 1A). After incubation with 10^−8^ M insulin for 24, 48, or 72 h, GH3 cells grow slower than untreated GH3 cells though without statistical significance (Figure 1B). The similar situation occurs in GH3 cells incubated with 10^−9^ M insulin for 24, 48, or 72 h (Figure 1C). These results unraveled 10^−7^ M insulin for 72 h incubation significantly represses tumor cell proliferation, and 10^−8^ or 10^−9^ M insulin does not enhance tumor cell proliferation.

**Figure 1.**
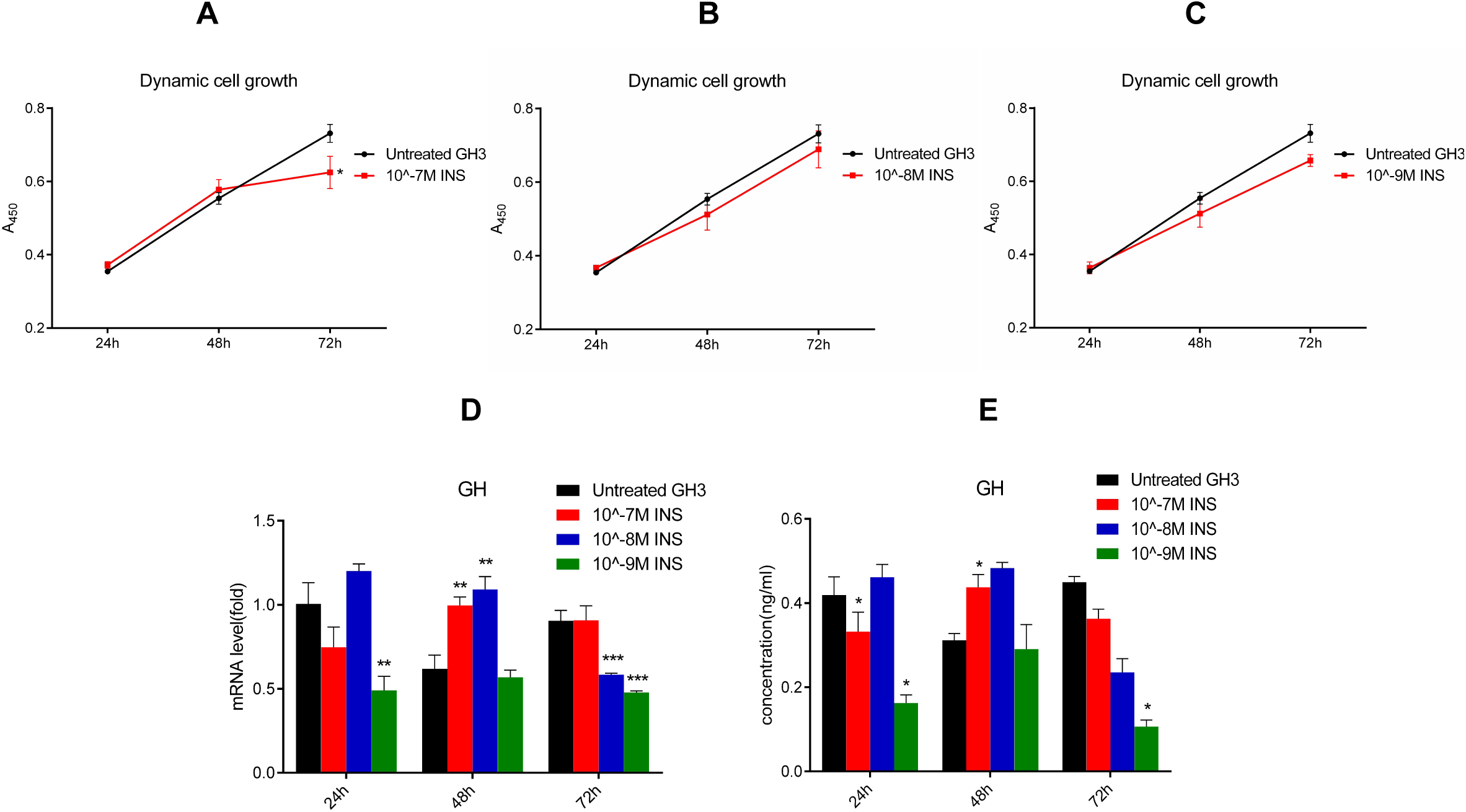
Effects of exogenous insulin on tumor cell proliferation and hormone secretion in GH3 cells grown in a high-glucose medium. (A-C) A_450_ values representing cell growth and measured by cell counting kit 8 (CCK-8) after incubation for 24, 48, or 72 h in 10^−7^ (A), 10^−8^ (B), or 10^−9^ M (C) insulin. (D) Fold changes of *GH* mRNA. (E) Concentrations of GH. Glucose concentration is 17.49 mM. Insulin dose is 10^−7^, 10^−8^, or 10^−9^ M, and incubation duration is 24, 48, or 72 h. INS: insulin. ∗: Significant difference from untreated GH3 cells (*p*<0.05, n=10). ∗∗: Very significant difference from untreated GH3 cells (*p*<0.01). ∗∗∗: Very very significant difference from untreated GH3 cells (*p*<0.001, *n*=10).

To reveal the implication of insulin in modulation of growth hormone (GH), we quantified its expressed mRNA and protein levels by quantitative polymerase chain reaction (qPCR) and Western blotting (WB). Consequently, it was clearly that untreated GH3 cells show a significantly higher level of *GH* mRNA than 10^−9^ M insulin-treated GH3 cells as measured on 24 h and 72 h (Figure 1D). Similarly, untreated GH3 cells also display a significantly higher level of GH than 10^−9^ M insulin-treated GH3 cells when tested on 24 h and 72 h (Figure 1E). Additionally, insulin remarkably downregulates *GH* mRNA and GH in 10^−9^ M for 72 h. These results of declined *GH* expression and diminished GH secretion might imply inhibitory tumor proliferation.

### Insulin-combined high-level glucose downregulates tumor marker gene expression

To more accurately evaluate the effect of insulin on tumor growth, we quantified the common tumor markers, cyclin 1 (CD1) and alpha fetoprotein (AFP), in the insulin-treated GH3 cells by qPCR and WB. After incubation with 10^−7^, 10^−8^, or 10^−9^ M insulin for 24, 48, or 72 h, GH3 cells show much lower levels of *CD1* mRNA (Figure 2A) and *AFP* mRNA (Figure 2B) than untreated GH3 cells with very significant statistical difference. From WB results illustrated in Figure 2C-2E, it was also noticeable that CD1 and AFP levels in 10^−7^, 10^−8^, or 10^−9^ M insulin-treated GH3 cells are almost lower than those in untreated GH3 cells after incubation for 24, 48, or 72 h. These results indicated downregulation of tumor marker expression is concomitant with suppression of tumor cell proliferation.

**Figure 2.**
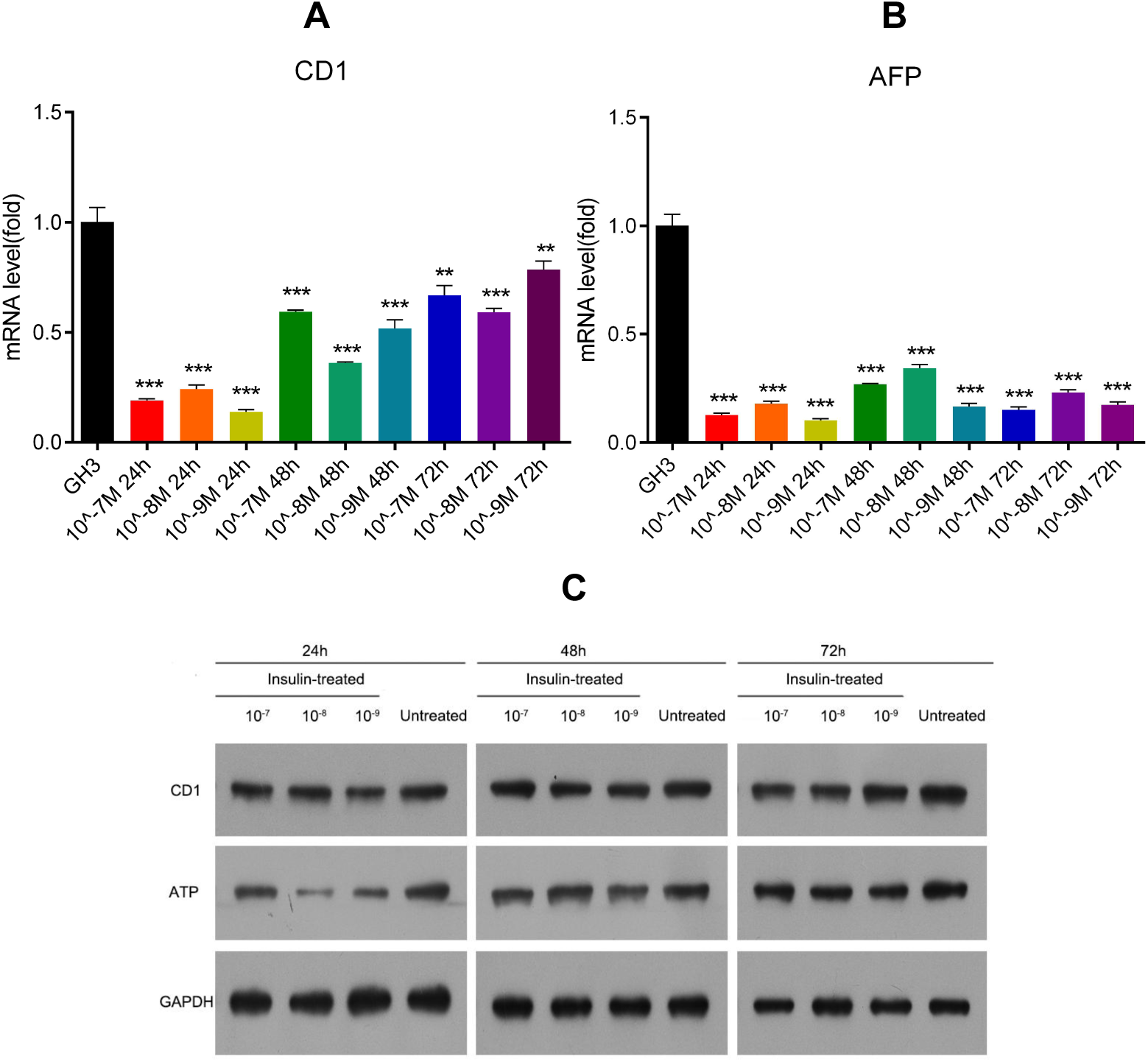
Effects of exogenous insulin on tumor marker gene expression in GH3 cells grown in a high-glucose medium. (A) Fold changes of *CD1* mRNA. (B) Fold changes of *AFP* mRNA. (C-E) Concentrations of CD1 and AFP. Glucose concentration is 17.49 mM. Insulin dose is 10^−7^, 10^−8^, or 10^−9^ M, and incubation duration is 24, 48, or 72 h. ∗∗: Very significant difference from untreated GH3 cells (*p*<0.01, n=10). ∗∗∗: Very very significant difference from untreated GH3 cells (*p*<0.001, *n*=10).

It was more remarkable from a comparison of relative gray-scale values estimated from CD1/glyceraldehyde-3-phosphate dehydrogenase (GAPDH) or AFP/GAPDH, in which 10^−9^ M insulin-treated GH3 cells for 24 h show the lowest CD1/GAPDH ratio, and 10^−8^ M insulin-treated GH3 cells for 24 h exhibit the lowest AFP/GAPDH ratio (Table 1). These results indicated insulin in the doses of 10^−7^, 10^−8^, or 10^−9^ M and during incubation for 24, 48, or 72 h declines CD1 and AFP levels at different extents. Although AFP was considered to be mainly synthesized in hepatic cells, it is also expressed in GH3 cells with an almost comparative level to CD1, perhaps representing a non-specific expression pattern.

**Table 1.** Comparison of CD1/GAPDH and AFP/GAPDH derived from the gray-scale values of WB bands in untreated GH3 cells and 10^−7^, 10^−8^, or 10^−9^ M insulin-treated GH3 cells for 24, 48, or 72 h

| Incubation time | Target protein/<br>Reference protein | $10^{-7}$ M insulin-treated GH3 | $10^{-8}$ M insulin-treated GH3 | $10^{-9}$ M insulin-treated GH3 | Untreated<br>GH3 |
| --- | --- | --- | --- | --- | --- |
| 24h | CD1/GAPDH | 0.64 | 0.58 | 0.35 | 0.61 |
|  | AFP/GAPDH | 0.35 | 0.05 | 0.14 | 0.50 |
| 48h | CD1/GAPDH | 0.82 | 0.68 | 0.64 | 0.81 |
|  | AFP/GAPDH | 0.37 | 0.52 | 0.33 | 0.59 |
| 72h | CD1/GAPDH | 0.74 | 0.65 | 1.12 | 1.31 |
|  | AFP/GAPDH | 0.85 | 0.78 | 0.80 | 1.06 |

### High-level glucose without insulin also downregulates tumor marker gene expression on the translational level

To simulate hyperglycemia occurring in DM patients, we employed a higher- glucose (30 mM) and insulin-free medium during GH3 cell culture. After incubation for 24, 48, or 72 h, it was obviously that *CD1* mRNA levels in 30 mM glucose-cultured GH3 cells are significantly higher as tested on 24 and 72 h, but equal to those in 17.49 mM-cultured GH3 cells on 48 h (Figure 3A), whereas *AFP* mRNA levels are significantly higher as tested on 48 and 72 h, but equal to those in 17.49 mM-cultured GH3 cells on 24 h (Figure 3B). Nevertheless, CD1 and AFP protein levels in 30 mM glucose-cultured GH3 cells are generally equal to those in 17.49 mM-cultured GH3 cells, only except for 72 h (Figure 3C). As tested on 72 h, for example, CD1/GAPDH and AFP/GAPDH is 0.41 and 0.18 in 30 mM glucose-cultured GH3 cells, whereas it is 0.05 and 0.13 in 17.49 mM glucose-cultured GH3 cells ( and Table 2). These transcriptional accelerated but translationally inhibited expression of *CD1* and *AFP* might be resulted from the general formation of a cellular starvation state because glucose uptake is proceeded slowly without insulin action.

**Figure 3.**
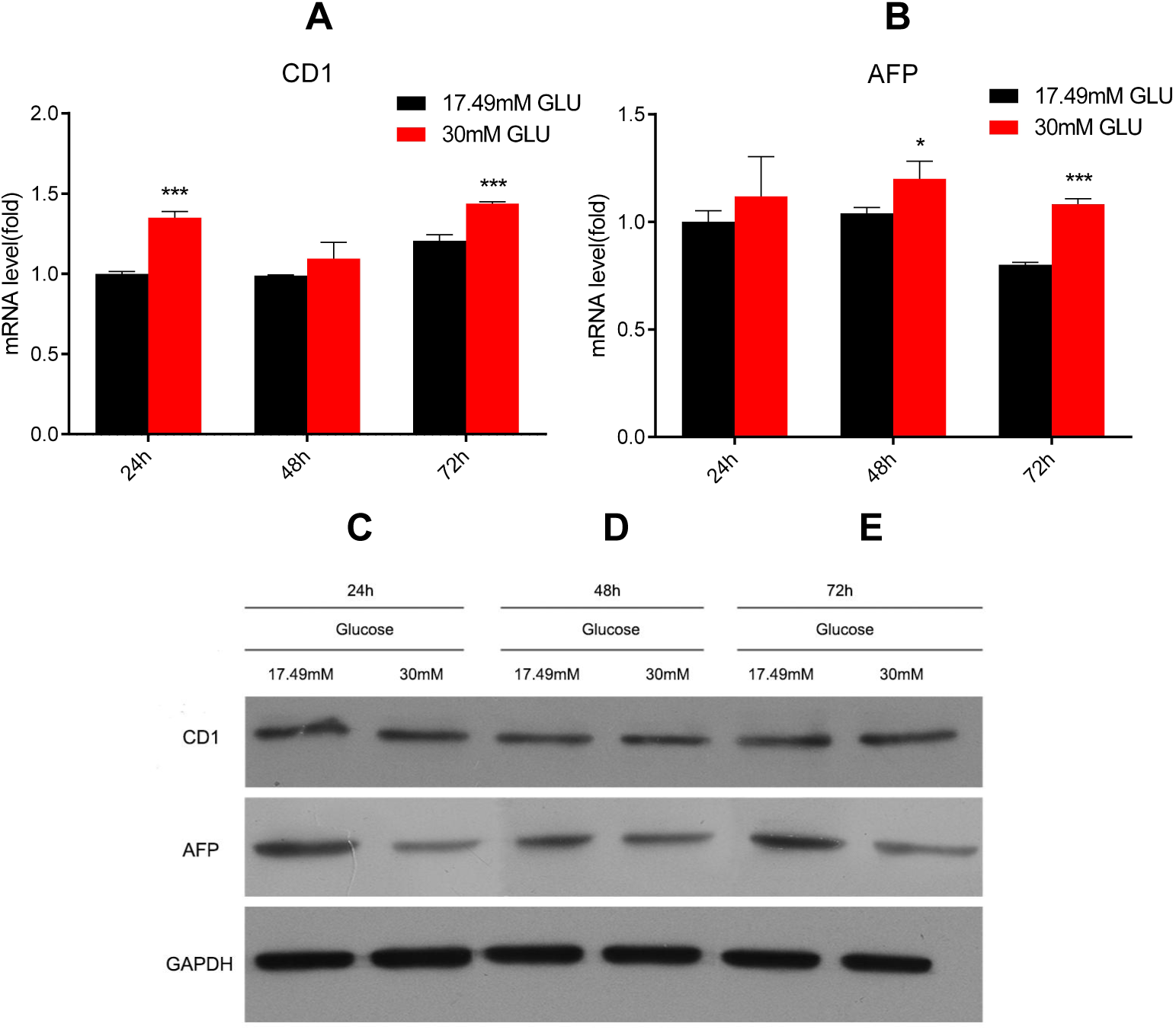
Effects of high-glucose on tumor marker gene expression in GH3 cells grown in insulin-free medium. (A) Fold changes of *CD1* mRNA. (B) Fold changes of *AFP* mRNA. (C-E) Concentrations of CD1 and AFP. Glucose concentration is 17.49 mM or 30 mM. Incubation duration is 24, 48, or 72 h. GLU: glucose. ∗: Significant difference from untreated GH3 cells (*p*<0.05, *n*=10). ∗∗∗: Very very significant difference from untreated GH3 cells (*p*<0.001, *n*=10).

**Table 2.** Comparison of CD1/GAPDH and AFP/GAPDH derived from the gray-scale values of WB bands in 17.49 mM glucose-cultured GH3 cells and 30 mM glucose-cultured GH3 cells for 24, 48, or 72 h

| Target protein/<br>Reference protein | 17.49 mM glucose-cultured GH3 |  |  | 30 mM glucose-cultured GH3 |  |  |
| --- | --- | --- | --- | --- | --- | --- |
|  | 24 h | 48 h | 72 h | 24 h | 48 h | 72 h |
| CD1/GAPDH | 0.34 | 0.27 | 0.05 | 0.25 | 0.24 | 0.41 |
| AFP/GAPDH | 0.45 | 0.25 | 0.13 | 0.09 | 0.17 | 0.18 |

### High-level glucose with or without insulin compromises ROS burst

To explore whether insulin might exert anti-oxidative roles, we determined the ROS levels in GH3 cells upon incubation with 10^−7^, 10^−8^, or 10^−9^ M insulin for 24, 48, or 72 h. From Figure 4A-4C, it was clearly seen that ROS levels in insulin-treated GH3 cells are much lower than those in untreated GH3 cells, in which the higher the insulin doses, the lower the ROS levels. For example, the fluorescence strengths representing ROS levels in 10^−7^, 10^−8^, or 10^−9^ M insulin-treated GH3 cells for 24 h are 59.3%, 81.0%, or 82.3%, whereas those in untreated GH3 cells were 96.0%, suggesting insulin might attenuate ROS burst or scavenge ROS already generated.

**Figure 4.**
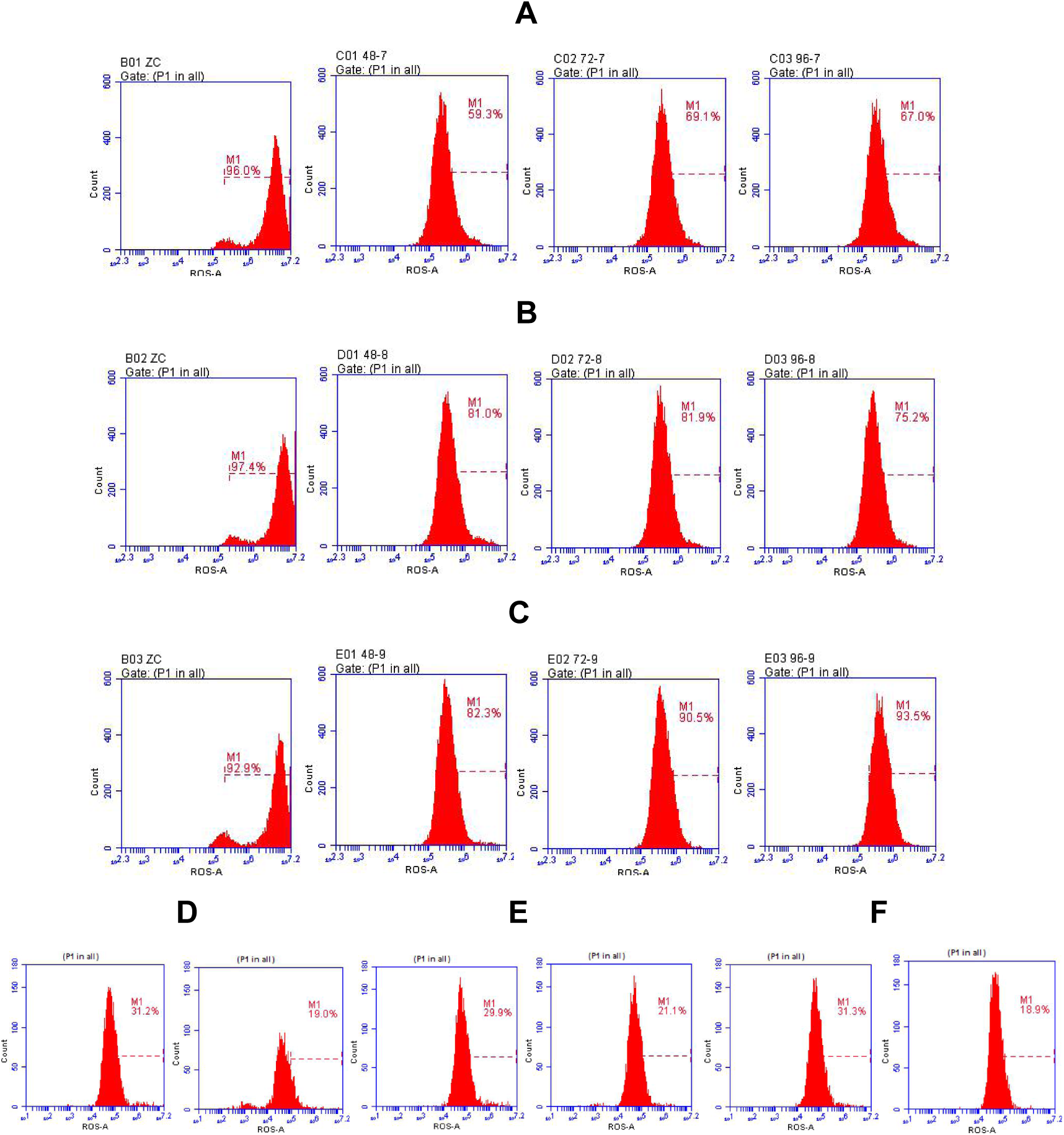
Effects of high-glucose combined with or without insulin on ROS production in GH3 cells. (A-C) ROS levels in 17.49 mM glucose-cultured GH3 cells after incubation with or without 10^−7^, 10^−8^, or 10^−9^ M insulin for 24 h (A), 48 h (B), or 72 h (C). (D-F) ROS levels in 17.49 mM or 30 mM glucose-cultured GH3 cells after incubation for 24 h (D), 48 h (E), or 72 h (F).

When incubation of GH3 cells with 30 mM glucose for 24, 48, or 72 h, ROS levels (Figure 4D-4F) are also unchanged or lower as compared with 17.49 mM glucose-cultured GH3 cells. For example, ROS levels are 21.1% and 29.9% in 30 mM glucose-cultured GH3 cells and 17.49 mM glucose-cultured GH3 cells after incubation for 72 h, respectively. These results suggested cellular starvation might compromise ROS burst.

### High-level glucose with or without insulin mitigates apoptosis

Following the decline of ROS levels, it was anticipated that ROS-induced apoptosis should be accordingly mitigated. As illustrated in Figure 5A-5C, it was shown that the total apoptotic cell percentages are extremely lower in 10^−7^, 10^−8^, or 10^−9^ M insulin-treated GH3 cells than those in untreated GH3 cells after incubation for 24, 48, or 72 h in the medium containing 17.94 mM glucose. For example, apoptotic cell percentages in 10^−7^, 10^−8^, or 10^−9^ M insulin-treated GH3 cells for 24 h are 20.5%, 20.0%, or 26.1%, whereas those in untreated GH3 cells are 32.6%, further addressing insulin enables compromise of ROS-driven apoptosis.

**Figure 5.**
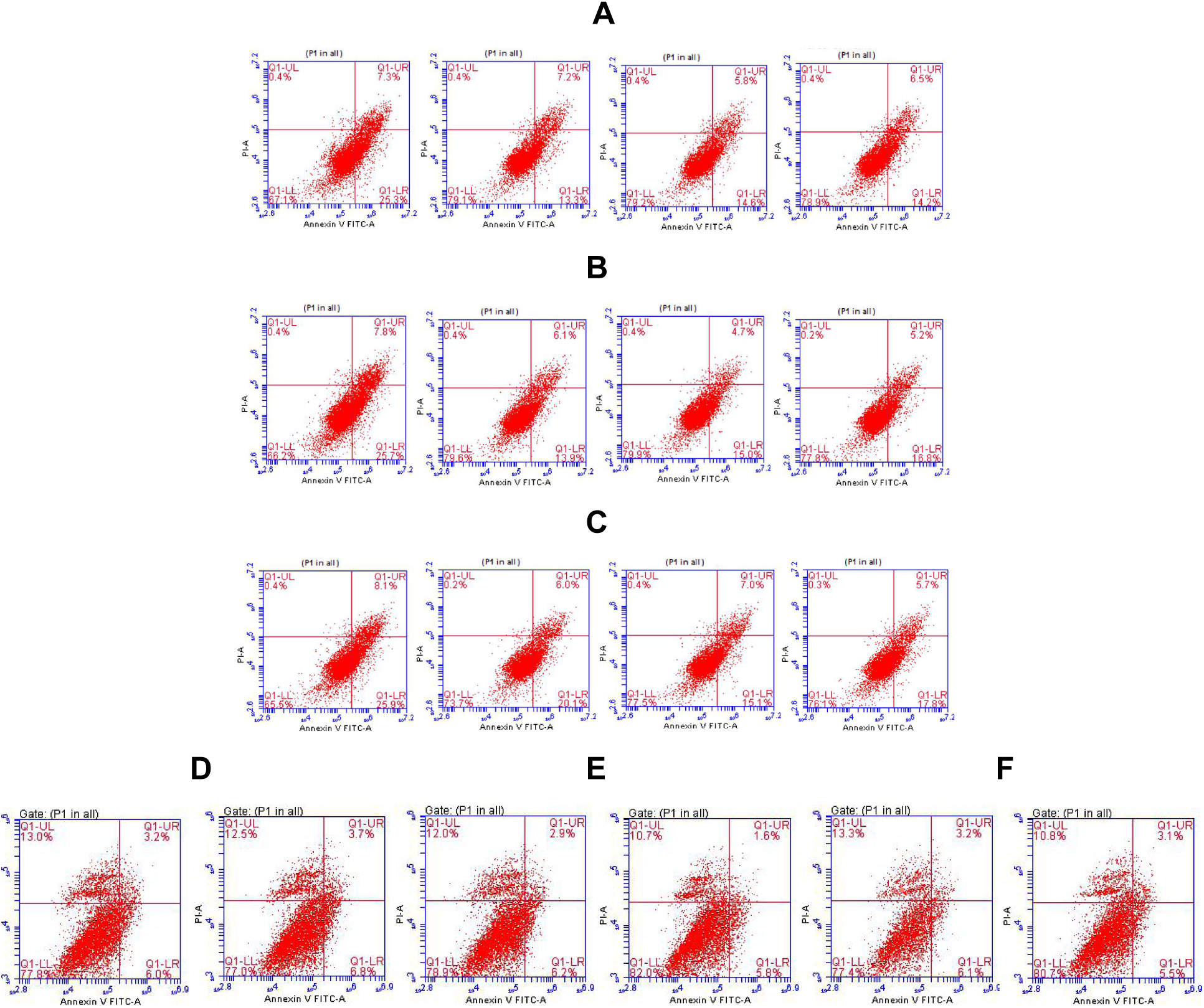
Effects of high-glucose combined with or without insulin on apoptotic induction in GH3 cells. (A-C) Apoptotic cell percentages in 17.49 mM glucose-cultured GH3 cells after incubation with or without 10^−7^, 10^−8^, or 10^−9^ M insulin for 24 h (A), 48 h (B), or 72 h (C). (D-F) Apoptotic cell percentages in 17.49 mM or 30 mM glucose-cultured GH3 cells incubated for 24 h (D), 48 h (E), or 72 h (F).

In similar, the late-phase apoptotic cell percentages after incubation for 24 h are almost equal (5.5% or 6.1%) in those cells treated with 17.94 mM or 30 mM glucose but without insulin (Figure 5D-5F). These results indicated that a high-level glucose but insulin-free medium weakens apoptosis induction, probably due to limited glucose uptake without insulin even under the glucose-rich environment.

### Knockdown of *NOS2* exhibits anti-tumor and anti-oxidation effects

To explore the possible mechanism underlying how can cellular starvation exert anti-tumor effects, we assumed cellular starvation might compromise the cell division responsible PI3K-Akt-mTOR signaling by downregulating insulin growth factor 1 (IGF-1). Thus, we quantified the expression level of *IGF*-*1* mRNA and IGF-1 in GH3 cells. As incubated with 10^−7^ M insulin for 24, 48, and 72 h, with 10^−8^ M insulin for 24 and 48 h, or with 10^−9^ M insulin for 24 and 48 h, *IGF*-*1* mRNA levels (Figure 6A) and IGF-1 levels (Figure 6B) are significantly lower than those in untreated GH3 cells. Intriguingly, GH3 cells even overexpress *IGF* mRNA and IGF after incubation with 10^−9^ M insulin for 48 h and 72 h. It was unable to be well explained at this moment, but it might be possible that compensation of an extremely low insulin level by IGF-1 overexpression. Because tumor cells are de-differentiated cells where genes are non-specifically expressed, IGF-1 was able to be detected in GH3 cells even though the pituitary gland is not an IGF-1-secreting organ.

**Figure 6.**
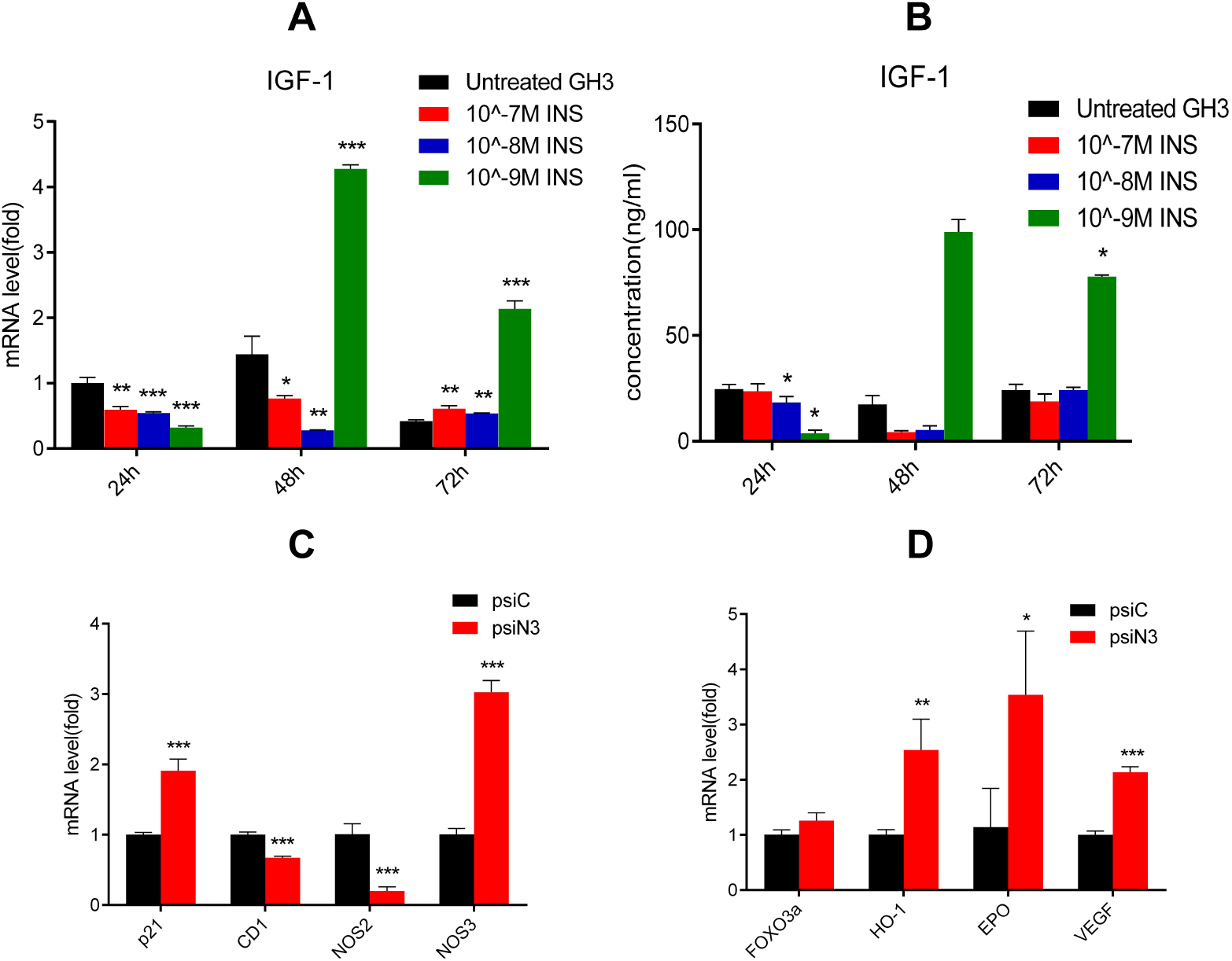
Effects of exogenous insulin on *IGF*-*1* expression in GH3 cells grown in a high-glucose medium (A and B) and effects of *NOS2* knockdown on the expression of anti-tumor and anti-oxidation responsible genes (C and D). (A) Fold changes of *IGF*-*1* mRNA. (B) Concentrations of IGF-1. Glucose concentration is 17.49 mM. Insulin dose is 10^−7^, 10^−8^, or 10^−9^ M, and incubation duration is 24, 48, or 72 h. INS: insulin. ∗: Significant difference from untreated GH3 cells (*p*<0.05, *n*=10). ∗∗: Very significant difference from untreated GH3 cells (*p*<0.01). ∗∗∗: Very very significant difference from untreated GH3 cells (*p*<0.001, *n*=10). (C) Fold changes of *p21*, *CD1*, *NOS2*, and *NOS3* mRNAs in HCC1937 (psiN3) cells. (D) Fold changes of *FOXO3a*, *HO*-*1*, *EPO*, and *VEGF* mRNAs HCC1937 (psiN3) cells. ∗: Significant difference from HCC1937 (psiN3) cells (*p*<0.05, *n*=6). ∗∗: Very significant difference from HCC1937 (psiN3) cells (*p*<0.01, *n*=6). ∗∗∗: Very very significant difference from HCC1937 (psiN3) cells (*p*<0.001, *n*=6).

To further explore the alternative mechanism behind cellular starvation exerting anti-tumor effects, we proposed CR-like anti-inflammatory responses might emerge as pivotal steps toward anti-tumor, which should downregulate *NOS2*-encoded inducible nitric oxide synthase (iNOS). Therefore, we investigated the putative relevance of *NOS2* knockdown with anti-oxidation and anti-tumor activity. By constructing *NOS2*-targeted small interfering RNA (siRNA) vectors and infecting them into breast cancer cells, we obtained the stable *NOS2* expression inhibitory cell line HCC1937 (psiN3) with the best interfering effect. From Figure 6C, it was evident that *NOS2*/iNOS downregulation is synchronous to upregulation of *NOS3*-encoded endothelial NOS (eNOS), accompanying with downregulation of the oncogene *CD1* and upregulation of the tumor suppressor gene *p21.* Furthermore, *NOS2* knockdown also upregulate the common anti-oxidation responsive genes, including the transcription factor *Forkhead box O3* (*FOXO3a*) that upregulates antioxidant enzymes, *HO*-*1* encoding heme oxygenase 1, *EPO* encoding erythropoietin, and *VEGF* encoding vascular endothelial growth factor (Figure 6D).

These results indicated *NOS2* knockdown can replicate CR-mediated anti-oxidant and anti-tumor effects, in which *CD1* downregulation was seen in CR-like cellular starvation, and upregulation of anti-oxidation responsible genes was also consistent with the outcome of compromised oxidative stress and ROS-induced apoptosis occurring in CR-like cellular starvation.

## Discussion

Upon incubation with different doses of insulin for different duration, GH3 cells exhibit growth inhibition with *IGF*-*1* downregulation, addressing insulin does not exert a tumor-promoting effect in a CR-resembling manner [21, 22]. Although IGF-1 is typically secreted by the liver upon accepting GH-conveying signals, it can also non-specifically expressed in tumor tissues with a de-differential feature. For example, melanoma-initiating cells can produce IGF-1 [23], and multiple myoloma cells can also generate IGF-1 [24]. Therefore, it could be acceptable that *IGF*-*1* downregulation can mimic metformin to repress tumor cell propagation because of impeded PI3K-Akt-mTOR signaling. An earlier study indicated metformin inactivates mTOR to inhibit breast cancer cell growth [25]. Clinically, metformin was observed to decrease the cancer rate for 25-40% via inactivating mTOR [26].

Whether insulin or high-glucose downregulates *IGF*-*1* remains unanswered at this moment, but we suppose herein that downregulation of *IGF*-*1* might be attributed to an intracellular low glucose milieu in tumor cells with insulin resistance. To replicate hyperglycemia seen in T1DM/T2DM, we cultured GH3 cells in the medium containing high-glucose (17.49 mM) or higher-glucose (30 mM) in combination with or without 10^−9^-10^−7^ M insulin. In other researches, 10, 15, or 20 mM glucose as well as 10^−7^ M insulin were separately exploited to induce insulin resistance in adipocytes and hepatoma cells [17, 18]. Regardless of insulin presence or absence, insulin resistance should be induced in GH3 cells by 17.49 mM or 30 mM glucose that resembles hyperglycemia. A high rate of hyperglycemia has been proven to be associated with inhibitors targeting IGF-1 receptor, which mediates a deleterious metabolic effect by decreasing insulin secretion and increasing insulin resistance in peripheral tissues [27]. Furthermore, a case of “pseudoacromegaly” has been reported to exhibit insulin resistance accompanying with a low IGF-1 level [28].

Laron syndrome (dwarfism), a genetic disease arisen from a mutant GH receptor gene, confers the low risks of DM and cancer [29]. The patients with Laron syndrome are characterized by a high level of GH (GH resistance) and a low level of IGF-1 [30]. Interestingly, we also observed a slow rate of cell growth is synchronous to a low level of IGF-1 in a high-glucose medium supplemented with insulin, suggesting an association of low-level IGF-1 with low-rate cell division [31]. Considering a dual downregulation of *IGF*-*1* and *NF*-*κB* by CR [21], and also because of NF-κB upregulate NOS2/iNOS, we postulated here that CR should exert anti-tumor effects via downregulating *NOS2*/iNOS. Indeed, we confirmed in the present study that *NOS2* knockdown not only decreases *NOS2*/iNOS and increases *NOS3*/eNOS, but also downregulates *CD1* and upregulates *p21*, suggesting CR-like anti-tumor activity is mediated by turning off iNOS and turning on eNOS.

On the other hand, we noticed decrease in ROS and reduction in apoptosis of GH3 cells grown in a high-glucose medium supplemented with insulin, which was also observed in CR [32]. In our previous work, CR in yeast and mice was also found to diminish ROS burst, compromise telomere shortening, and promote lifespan extension [33, 34]. It is well-known that resveratrol is an activator of SIRT, metformin is an activator of AMPK, and CR activates AMPK, SIRT1, and PGC-1α [35, 36]. PGC-1α in turn promotes mitochondrial biogenesis for multiple beneficial effects [37]. Therefore, it was understood that CR-like anti-oxidant effects should mitigate ROS burst and attenuate ROS-induced apoptosis. In the experimental confirmation of CR-like anti-oxidant effects by *NOS2* knockdown, we noticed the significant upregulation of a series of anti-oxidation responsible genes. FOXO3a serves as a tumor suppressor for protection from oxidative stress by upregulating antioxidant enzymes such as catalase and superoxide dismutase (SOD) [38]. HO-1 reduces oxidative stress by degrading the pro-oxidant heme [39]. EPO and VEGF promote erythropoiesis and angiogenesis to attenuate anemia and hypoxia [40, 41].

Intriguingly, we found treatment of GH3 cells by high-level glucose without insulin even gives rise to the similar consequence as treatment of GH3 cells by high-level glucose with insulin. We assumed the high-level glucose might downregulate glucose transporters and decreases glucose intake. A previous work showed sustained hyperglycemia *in vitro* downregulates GLUT1 [16]. However, an *in vitro* hyperglycemic environment was demonstrated to promote tumor progression by upregulation of GLUT1 and GLUT3 [42]. A five-year clinic observation also indicated hyperglycemia is highly related to aggravation of rectal tumors [43]. These contradictory outcomes would be reasonably interpreted by discriminating insulin sensitivity from insulin resistance. On one hand, insulin deficiency-caused hyperglycemia in T1DM with insulin sensitivity might show cancer promotion once insulin was administered and glucose uptake accelerated, which is also the reason why insulin is effective for hypoglycemia in T1MD patients. On the other hand, insulin receptors (IR) dysfunction-originated hyperglycemia in T2MD with insulin resistance might exhibit cancer prohibition regardless insulin was administered or not, which is also the reason why insulin is ineffective and metformin was effectively used in T2MD patients.

Alternatively, the above inconsistent data could be also explained by inactivation of PTEN due to the germ-line gene mutation [44] or suppression of PTEN transcription by NF-κB directly binding to the PTEN promoter under a pro-inflammatory circumstance [45]. This is because PI3K-activated glucose transport into cells via GLUT is antagonized by PTEN [46]. When PI3K is over-activated, cells prone to intake more glucose for anaerobic respiration (Warburg effects), which accounts for 10 to 20-fold increases of metabolic activity that predisposes mutagenesis [47]. These results implied PTEN should be in deficiency under the inflammation niches and hence some T1MD/T2MD patients might exhibit enhanced PI3K-Akt-mTOR signaling toward higher cancer risks.

Conclusively, insulin neither enhances pituitary adenoma cell propagation nor upregulates *CD1* and *AFP* expression in 17.49 mM or 30 mM glucose medium supplemented with or without 10^−7^-10^−9^ M insulin during 24-72 h of incubation. Mechanically, a high-glucose environment that induces insulin resistance and blocks glucose transport can mimic CR to exert anti-tumor effects by dually downregulating *IGF*-*1* and *NOS2* expression. Whether insulin use would increase the cancer risk in the subjects with the insulin sensitive early-phase DM merits further clinical elucidation.

## Materials and Methods

### Cell culture and insulin incubation

The rat pituitary adenoma cell line GH3 was cultured in 25 cm^2^ cultural vessels and placed in a humidified incubator at 37°C with an atmosphere of 5% CO_2_ and 95% O_2_. Unless indicated, cells were cultivated in a 17.49 mM *D*-glucose-containing DMEM/F12 liquid medium (Hyclone, Cat. No. SH30023.01B) supplemented with heat-inactivated fetal bovine serum (Gibco, Cat. No. 10099-141), 100 U/ml penicillin, and 100 g/ml streptomycin. Alternatively, cell were cultivated in a 30 mM *D*-glucose-containing DMEM liquid medium (Hyclone, Cat. No. SH30022.01B, containing 24.51 mM *D*-glucose) supplemented with 5.49 mM *D*-glucose and heat-inactivated fetal calf serum (Gibco, Cat. No. 10099-141), 100 U/ml penicillin, and 100 g/ml streptomycin was also included. Upon centrifuged at 1000 r/min for 5 min, cell suspensions were transferred into 6-well culture plates for incubation in a 17.49 mM glucose-containing DMEM/F12 liquid medium with human insulin (Sigma, Cat. No. I3536) in a dose of 10^−7^, 10^−8^, or 10^−9^ M for 24, 48, or 72 h, or in a 30 mM glucose-containing DMEM liquid medium for 24, 48, or 72 h. After completion of culture, cells were pooled by centrifugation for further analysis.

### qPCR

RNA isolation, quality control, electrophoresis, reverse transcription, and quantification were performed using Agilent Stratagene MX3000P in a PCR reaction (92°C 2 min, 94°C 20 sec, 58°C 20 sec, 72°C 20 sec, 40 cycles) by following forward (F) and reverse (R) primers designed for the reference gene *β*-*actin* and all target genes.

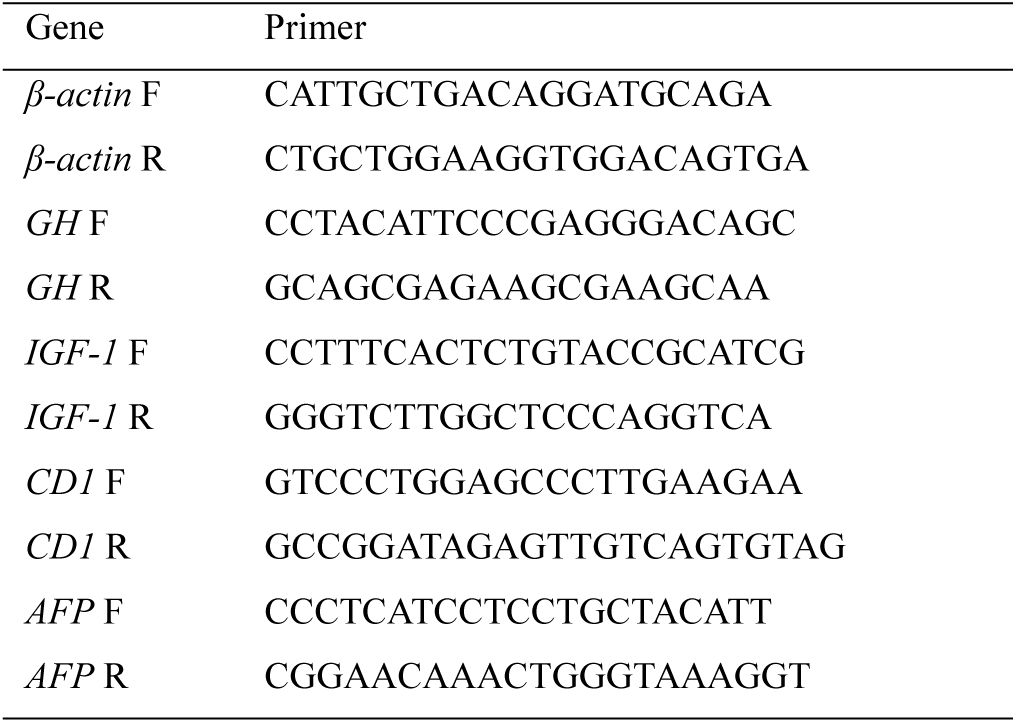

The copy numbers of amplified genes were estimated by 2^−ΔCT^, in which ΔCT = (target gene (treatment group) / target gene (control group)) / (housekeeping gene (treatment group) / house-keeping gene (control group)). The raw qPCR data were normalized by the copy numbers of the reference gene *β*-*actin*, and the fold changes of each target gene were calculated by comparing the 2^-ΔCT^ value of a treatment sample with that of a control sample.

### WB and enzyme-linked immunosorbent assay (ELISA)

The reference protein GAPDH and all tested antigen proteins were immunoquantified according to manufacture’s manuals. Anti-CD1 antibody (H-295, Cat. No. sc-753) and anti-AFP antibody (C-19, Cat. No. sc-8108) were purchased from Santa Cruz Biotechnology Inc., USA. The ELISA Kits for rat GH (Cat. No. F3997-A) and rat IGF-1 (Cat. No. F8278-A) were purchased from Shanghai Kexing Biotechnology Ltd Co., China.

### Dynamic cell growth monitoring

After staining cells with CCK-8, absorbance at the wave length of 450 nm (A_450_) was measured using a microplate reader (Thermo Fisher Scientific, Multiscan MK3) to monitor dynamic cell growth.

### ROS and apoptosis analysis

ROS Assay Kit (S0033) was obtained from Beyotime Institute of Biotechnology, China. Annexin V/PI Apoptosis Kit (AP101) and Cell Cycle Staining Kit (CCS012) were supplied by Hangzhou MultiScience (Lianke) Biotech Co., Ltd., China. ROS and apoptosis were measured by a flow cytometer (BD, FACS Calibur) using corresponding reagent kits and based on manufacturer’s instructions.

### *NOS2* knockdown by short-hairpin RNA interference

By inserting one of three candidate siRNA fragments, siRNA-*NOS2*-1/2/3 (abbrev. siN1/2/3, interfering with one of three selected *NOS2* target sequences into the expression plasmid vector pSUPERretro, three recombinant siRNA-*NOS2*-expressing constructs, pSUPERretro-siN1/2/3 (abbrev. psiN1/2/3), were obtained. After transiently infecting one of them into HCC1937 cells, *NOS2* mRNA and iNOS levels were quantified in HCC1937 (psiN1/2/3) cells. Among the transformed cells, NC (psiN3) cells with the lowest *NOS2* mRNA and iNOS levels were obtained. So psiN3 was chosen to stably transfect HCC1937 cells for establishing a stable cell line HCC1937 (psiN3).

### Statistical analysis

The software SPSS 22.0 was employed to analyze data, and the software GraphPad Prism 5.0 was employed to plot graphs. The Independent Simple Test was used to compare all groups, but the Kruskal-Wallis Test followed by Nemenyi test was used when the data distribution is skewed. The significance level (*p* value) was set at <0.05 (∗), <0.01 (∗∗), and <0.001 (∗∗∗).

## Acknowledgments

We thank our colleagues in Tropical Medicine Institute of Guangzhou University of Chinese Medicine. This work was supported by the National Natural Science Foundation of China (No. 81273620 to Q.P.Zeng and No. 81673861 to C.Q. Li) and by the High-Level University Construction Foundation of Guangdong Province of China (No. 2050205).

## Author Contributions

Conceptualization: QPZ and CQL. Data curation: QPZ and TL. Formal analysis: TL; XAH. Funding acquisition: QPZ; CQL; XHL. Investigation: TL; YPC; LLT. Methodology: TL; JDZ. Project administration: SQH; QX. Resources: QX; XAH. Software: XAH. Supervision: QX. Validation: QPZ; QX. Visualization: QPZ; TL. Writing original draft: QPZ; CQL. Writing review & editing: QPZ; CQL; TL.

